# Random transposon mutagenesis identifies genes essential for transformation in naturally competent archaea

**DOI:** 10.1101/2022.08.17.504363

**Authors:** Dallas R. Fonseca, Madison B. Loppnow, Leslie A. Day, Elisa L. Kelsey, Mohd Farid Abdul Halim, Kyle C. Costa

**Affiliations:** Department of Plant and Microbial Biology, University of Minnesota, Twin Cities, St Paul, MN

## Abstract

Natural transformation, the process whereby a cell acquires DNA directly from the environment, is an important driver of evolution in microbial populations. While transformation is well characterized in bacteria, relatively little is known about this process in archaea. Here, we leverage an optimized method to generate transposon mutants in *Methanococcus maripaludis* to screen for genes essential to natural transformation. A screen of 5,376 mutant strains identified 25 candidate genes. Among these are genes encoding components of the type IV-like pilus, transcription/translation associated genes, putative membrane bound transport proteins, and genes of unknown function. Interestingly, similar genes were identified regardless of whether replicating or integrating plasmids were provided as substrate for transformation. Using allelic replacement mutagenesis, we confirmed that several genes identified in these screens are essential for transformation. Finally, we identified a homolog of a membrane bound substrate transporter in *Methanoculleus thermophilus* and verified its importance using allelic replacement mutagenesis, suggesting a conserved mechanism for DNA transfer in multiple archaea. These data provide an initial catalog of genes important for transformation in the archaea and can inform efforts to understand gene flow in this domain.

**Importance:** Horizontal gene transfer (HGT) is an important driver of evolution in microbial populations. One of the primary ways microorganisms acquire genetic material through HGT is transformation, the direct uptake of DNA from the environment. While transformation is well-studied in bacteria, little is known about this process in archaea. Using a random mutagenesis screen to identify components of the archaeal transformation pathway, we identify a catalog of genes important to transformation in *Methanococcus maripaludis* and show that a subset of these genes is functionally conserved across diverse archaea. This is a key step in understanding mechanisms of gene flow in natural populations, and identification of the DNA uptake system will assist in establishing new model genetic systems for studying the archaea.

## Introduction

Horizontal gene transfer, the process by which genetic information is transferred independent of inheritance, is a major driver of evolution in microbial populations. This typically occurs via one of three mechanisms: conjugation, the uptake of DNA via cell-to-cell contact; transduction, DNA transfer mediated via a virus/phage; or transformation, the direct uptake of DNA from the environment. In the archaea, mechanisms of transduction have been described and conjugation-like systems are known (1–5); however, little is known about natural transformation-mediated DNA uptake.

Transformation has been well studied in numerous bacteria such as *Vibrio* spp., *Neisseria* spp., and *Bacillus* spp. (6–11). To enable transformation, a cell enters a physiological state known as the competent state, which can be achieved by sensing cues from other organisms, quorum sensing, starvation, or other environmental stressors (12). Some organisms (e.g. *Helicobacter pylori*) are hypothesized to be constitutively competent, but external stimuli can alter transformation efficiency in these organisms (6, 13). In the naturally competent archaeon *Methanococcus maripaludis* strain JJ, there was no difference in transformation frequency between exponential phase cell and stationary cells, suggesting that the strain may be constitutively competent (14). Interestingly, *M. maripaludis* strain S2 is not natively competent in our laboratory conditions; however, it can be induced into the competent state by expression of pilin proteins from a heterologous expression vector (14), suggesting that there may regulatory control of competence in *M. maripaludis*.

Once a cell has entered a competent state, transformation begins with the localization of free DNA to the cell surface (in the case of gram-negative bacteria, free DNA is transferred across the outer membrane into the periplasm). DNA localization is facilitated by extracellular appendages such as type IV pili (8–11, 15), the Flp pilus (16), the competence pilus (7), etc. In *M. maripaludis* and *Methanoculleus thermophilus*, a type IV-like pilus is essential for natural transformation (14). We hypothesize that archaeal type IV-like pili and bacterial pili play similar roles in transformation.

After binding by pili, DNA enters the cell. In bacteria, double stranded DNA (dsDNA) is converted to single stranded DNA (ssDNA) before transport by a DNA transporter such as ComEC (6, 17). The DNA transporter in naturally competent archaea is unknown, and homologs of ComEC have not been identified in these organisms (18). Additionally, it is unclear whether dsDNA or ssDNA is the substrate for transformation in archaea. The final step of transformation is the incorporation of DNA into the genome by ssDNA binding proteins and recombinases such as RecA (19, 20).

In order to better understand the process of transformation in naturally competent archaea, we sought to determine which genes are essential for transformation. By employing a method for reproducible and high-efficiency transposon mutagenesis in *M. maripaludis*, we developed a screen for identifying defects in natural transformation. From 5,376 transformants screened, we identified 71 with insertions corresponding to 25 genes that resulted in a defect in transformation, representing ∼1.5% of the 1,815 genes encoded on the *M. maripaludis* strain JJ genome (21). Markerless gene deletion was used to confirm the transformation defect phenotype of these genes. Interestingly, we found an essential role for a membrane bound TctA-like protein for transformation in two distinct organisms. The TctA-like protein belongs to a family of transporters, but its function has not been characterized in archaea.

## Results

### Optimizing *M. maripaludis* mini-mariner transposon mutagenesis

To identify genes important for natural transformation in *M. maripaludis*, we employed random transposon mutagenesis to screen for mutants with a transformation defect. In our hands, previous methods for generating transposon mutants were either low efficiency, labor intensive, or irreproducible (22–24). Thus, we sought to optimize the methods of Sattler *et al*. and Quitzke *et al*. (22, 23) to improve efficiency. Using HimarI transposase and a mini-mariner transposon (25), we incubated transposon DNA with at least a 2-fold stoichiometric excess of purified transposase before transfer into *M. maripaludis* via the polyethylene glycol (PEG) method of transformation (26). Notably, the protocol we employed differed from the established method in that excess transposase was included in the reaction mixture, incubation time/temperature was optimized before transformation, and a heating step/DNA precipitation that likely inactivated the transposase was omitted. Following the optimized protocol, we observed a transformation efficiency averaging 5,367 colony forming units (CFUs) (µg DNA)^-1^ (Figure 1A), over 1000-fold more efficient than other published protocols.

**Figure 1.**
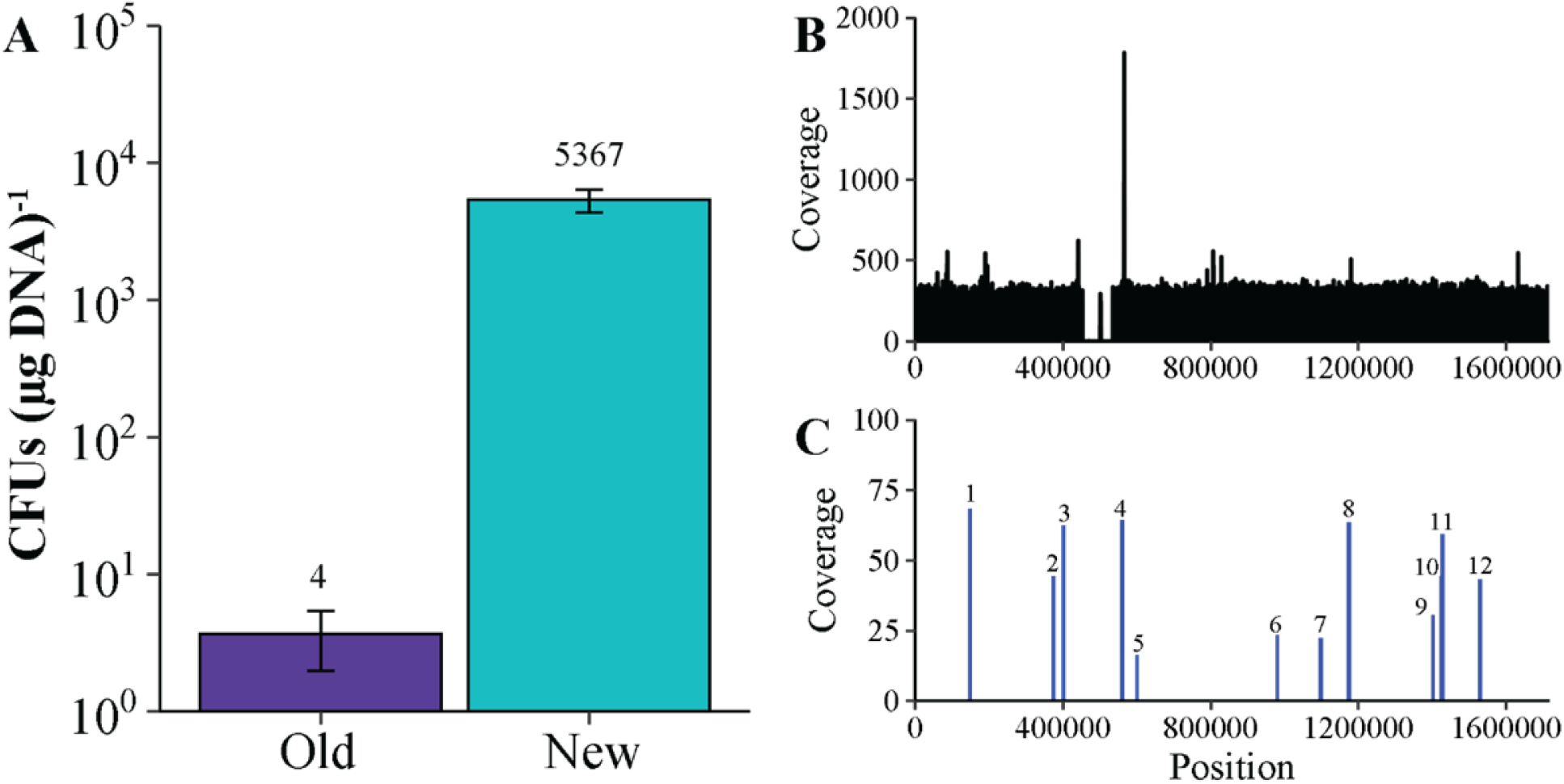
Optimized transposon mutagenesis produces random insertions across genome that can be mapped with whole genome sequencing: (A) Comparison of methods described by Sattler *et al*. (22) and Quitzke *et al*. (23) [old] and the method described in this study [new]. Data are averages from three independent experiments, and error bars represent one standard deviation around the mean. (B) Coverage distribution of reads mapped from a representative sequencing run. Coverage data created by Bowtie2 alignment of all forward reads to reference genome of *M. maripaludis* JJ (CP026606). (C) Sites of random transposon insertion across genome. Two Illumina reactions containing 6 random mutants each were sequenced and mapped to identify mini-mariner transposition. Insertion sites position and coverage value were determined through visualization in Geneious prime^®^(v.2021.0.3). Site determined by presence of AT (known integration site for mariner-based transposition) and where reads of either end of the transposon overlapped.

To validate that each CFU observed arose from a single transposon insertion and that transposon insertion sites were random, we mapped the insertions from 12 mutants across two sequencing reactions. Each sequencing reaction had relatively uniform coverage across the genome (Figure 1B). Additionally, mapping reads that aligned to both the genome and to the transposon identified 12 independent insertion sites (Figure 1C), validating our improved method for generating random transposon insertion mutants. Interestingly, while mapping via whole genome sequencing, we found that the genome of our strain of *M. maripaludis* strain JJ differed from the published genome (21) which in that it has an apparent ∼80 kbp deletion (Figure 1B).

However, we note that our strain retains the natural transformation phenotype despite this difference, so it was not considered further.

### Developing a screen for transformation defects with replicating plasmids

Using transposon mutagenesis, we developed a screen to identify genes important to transformation in *M. maripaludis*. 2,496 individual mutants were grown in 96 well plates to stationary phase in the absence of selection. After growth, wells were supplemented with 1 µg mL^-1^ of the replicating plasmid pLW40neo (27) and incubated for an additional day to allow for DNA uptake and expression of the antibiotic resistant cassette (14). Cultures were transferred in duplicate into medium containing neomycin and allowed to grow for 2 days, consistent with the time needed for selection of *M. maripaludis* transformants in liquid medium (14). Any mutants that failed to grow in both wells containing neomycin were considered candidates for a transformation defect. For further validation, candidate mutants were grown in 5 mL liquid culture and rescreened for a transformation defect (14). Cultures with an OD_600_ > 0.2 at 48 hours were considered false positives and were not analyzed further. In total, 46 mutants were selected for further analysis.

Transposon insertion sites were mapped to the *M. maripaludis* JJ reference genome (21). Of the 46 mutants sequenced, we identified insertions that likely impacted expression of 21 genes (Figure 2, Table 1). The majority of mutants had transposon insertions in genes encoding components of the type IV-like pilus, consistent with the importance of pili in natural transformation (14) and validated that our screen was revealing genes relevant to natural transformation. We additionally identified genes that encoded proteins with at least one predicted transmembrane helix (MMJJ_13020 and MMJJ_07810/7800) and therefore may be performing their function in/around the membrane. Of the remaining genes identified, some were predicted to be or be in putative operons with DNA/RNA binding proteins, possibly involved in transcription or translation (MMJJ_01810, MMJJ_13210, MMJJ_17710, and MMJJ_14410). Lastly, we identified multiple insertions in a 2 gene operon composed of a predicted MinD/ParA ATPase (MMJJ_11080) and a hypothetical protein (MMJJ_11090).

**Table 1.**
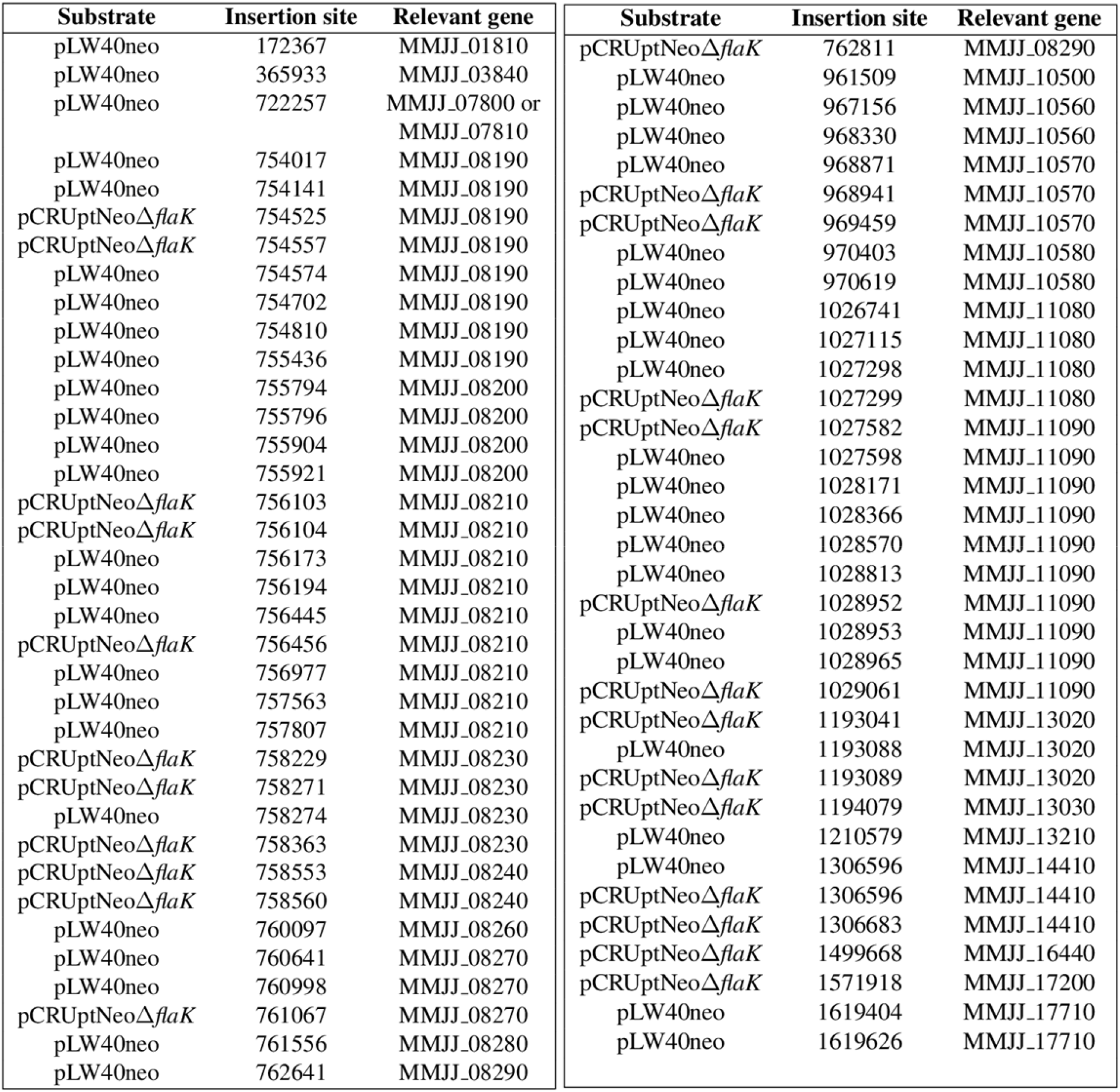
Insertions identified across both transposon screens.

**Figure 2.**
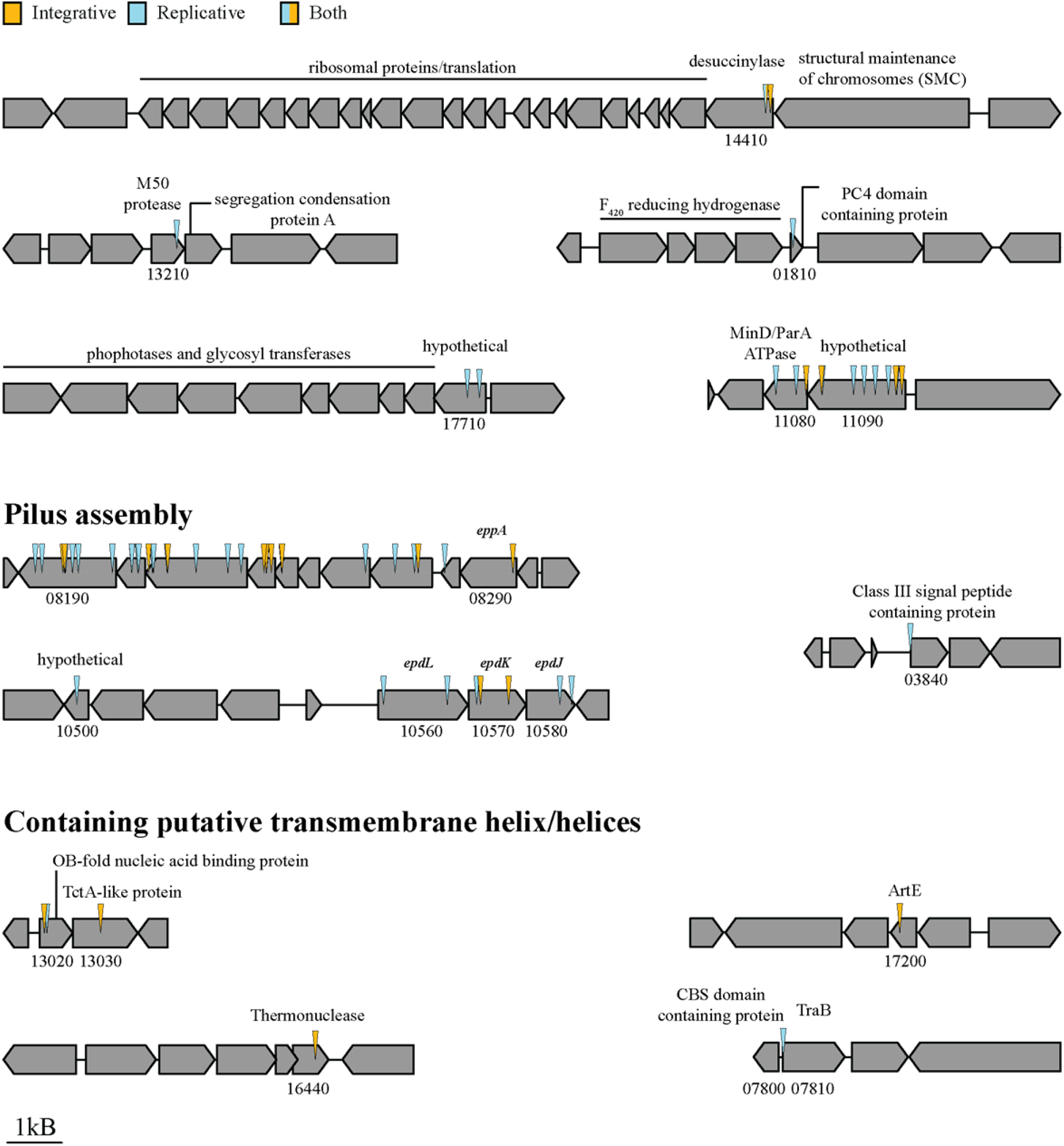
Schematic representation of insertions and respective operons: For each candidate mutant, insertion site was determined via visualization in Geneious prime^®^(v.2021.0.3). Site determined by presence of AT (known integration site for mariner-based transposition) and where reads of either end of the transposon overlapped. Genes around the insertion are included for context. Genes of interest are referenced using the locus tag format of Poehlein *et. al* (MMJJ_XXXXX)(21). Select annotations were added manually based on either NCBI or UniProt annotations. Colors used to mark which DNA substrate used in screen where replicative refers to pLW40neo and integrative refers to pCRUptNeoΔ*flaK*. One insertion in MMJJ_14400 was identified in both substrates, therefore it was colored with both. A table summary for each of these insertions is provided as table 1.

### Developing a screen for transformation defects with integrating plasmids

There is a significant difference between the transformation efficiencies of replicative versus integrative plasmids in *M. maripaludis*. While it was hypothesized that a requirement for recombination of integrating vectors into the genome accounted for this difference (14), it was recently reported that the presence of PstI restriction sites (5’-CTGCAG-3’) on a plasmid alters transformation efficiency (28). The integrating vector pCRUptNeo is typically used for mutagenesis of *M. maripaludis* and contains several PstI restriction sites. This is consistent with the presence of a PstI-like restriction system in *M. maripaludis* (26). Previously, it was suggested that MMJJ_06980, a predicted type-II restriction enzyme annotated as AplI (an isoschizomer of PstI) was responsible for the observed PstI activity in transformed cultures (21). We hypothesized that deletion of MMJJ_06980 would result in higher transformation efficiencies for integrating plasmids. A mutant strain lacking MMJJ_06980 was generated and transformed with the integrating vector pCRUptNeoΔ*flaK* (14). Transformation of the *M. maripaludis* Δ*MMJJ_06980* strain was 10-fold more efficient than transformation of wild type (Figure 3) suggesting that the native *M. maripaludis* PstI restriction activity was eliminated in the mutant. There was not a significant difference in transformation efficiency between wild type and *M. maripaludis* Δ*MMJJ_06980* when using replicating plasmid pLW40neo (Figure 3).

**Figure 3.**
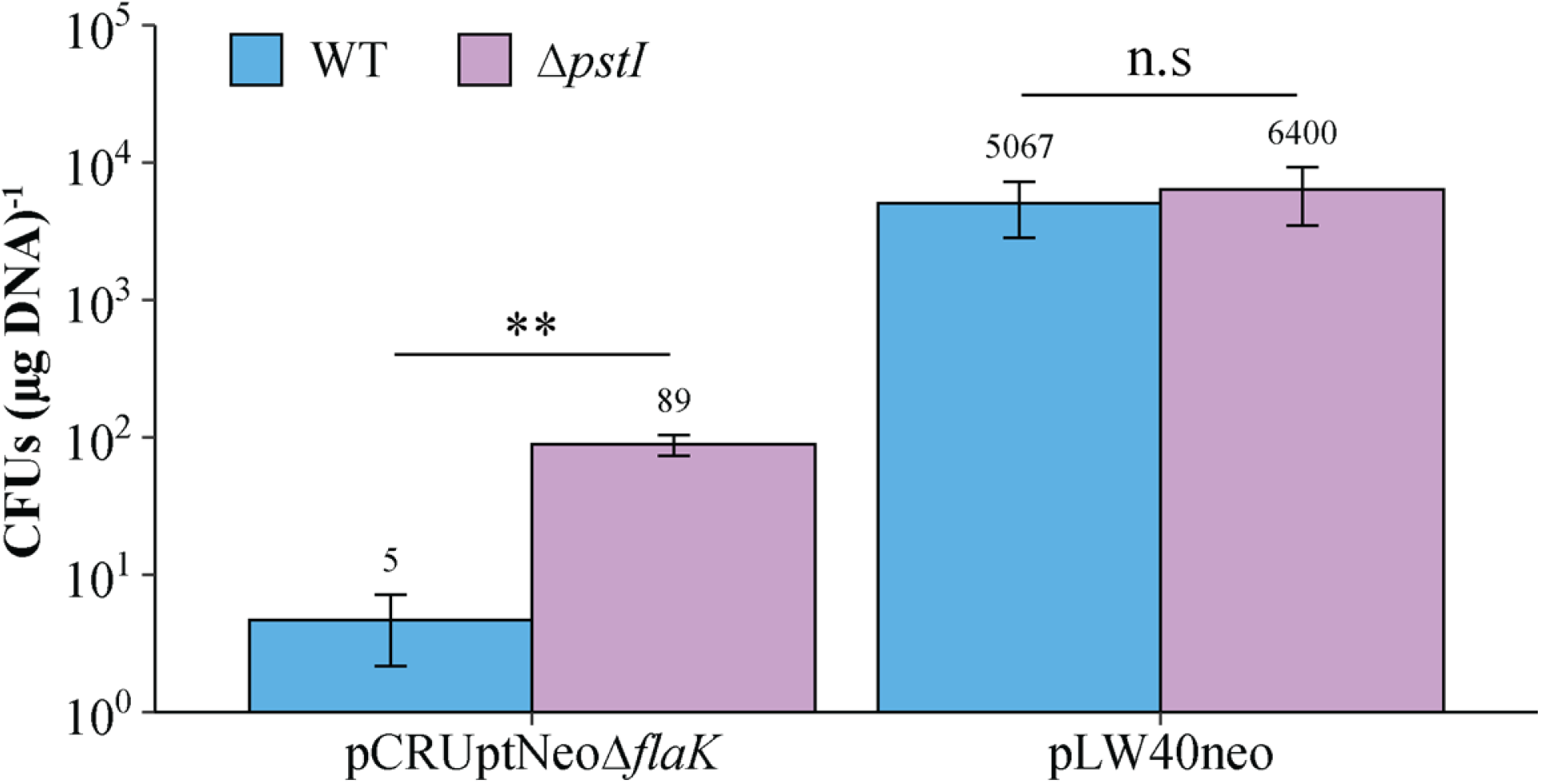
Deletion of MMJJ_06890, a predicted PstI restriction enzyme, increases the transformation efficiency of transformations with integrative plasmids: Transformation efficiencies of WT compared to Δ*MMJJ_06890* with either pCRUptNeoΔ*flaK* or pLW40neo. Data are averages from three independent experiments, and error bars represent one standard deviation around the mean. Statistics were performed using a two-tailed, equal variance T test. *, P < 0.05; **, P < 0.01; ***, P < 0.001

The increased transformation efficiency of the Δ*MMJJ_06980* strain allowed us to screen pCRUptNeoΔ*flaK* for transformation defects to determine if a different set of genes is essential for transformation with integrating vectors. We screened 2,880 transposon mutant strains and identified 25 with defects for transformation affecting 14 genes (Figure 2). Most of the transposon insertions fell into either the same genes or genes in putative operons with those identified in the pLW40neo-based screen (such as MMJJ_13030), suggesting that the pathways transformation are similar regardless of substrate. We additionally obtained transposon insertions in additional membrane associated proteins of unknown function, namely MMJJ_16440 (predicted thermonuclease) and MMJJ_17200 (predicted archaeosortase homolog, *artE*).

### MinD/ParA-like proteins and a putative cytoplasmic thermonuclease are needed for transformation

To validate both transposon mutagenesis screens, we constructed in-frame deletion mutants for several of the genes identified and tested their ability to take up exogenous DNA. We did not generate in-frame deletion mutants for genes encoding components of the type IV-like pilus as previous work already identified several of these as essential for transformation (14). While many of these genes were identified from both screens, all in-frame deletion mutants were tested for transformation defects with pLW40neo as the substrate.

MMJJ_11080 is a predicted MinD/ParA-like protein. Proteins from this family may function in polar localization of macromolecules. As such, they may be important for DNA segregation to daughter cells or localizing protein complexes to the cell pole (29). In a putative operon with this gene is MMJJ_11090, which encodes a hypothetical protein. In-frame deletion mutants of either gene were completely defective for transformation, and *in trans* expression of each gene in the respective deletion mutans rescued this transformation defect (Figure 3).

MMJJ_16440 is predicted to encode a thermonuclease with a secretion signal at the N-terminus. MMJJ_17200 is predicted to encode ArtE, an uncharacterized member of the archaeosortase protein family (30) where a known member is involved in C-terminal anchoring and processing of archaeal surface proteins (31). In-frame deletion mutants for each of these genes were completely defective for transformation, and complementation of the mutation *in trans* rescued this defect (Figure 3).

One candidate mutant with a defect in transformation contained a transposon insertion near the 5’ end of MMJJ_07810, a gene encoding a putative TraB-like protein. However, a Δ*MMJJ_07810* mutant did not have a defect in natural transformation efficiency. We hypothesized that the close proximity of this insertion to the promoter of MMJJ_07800 may have resulted in the observed transformation defect. However, a Δ*MMJJ_07800* deletion strain was also capable of DNA uptake (Figure 3). It may be that another mutation elsewhere on the genome of the transposon insertion strain resulted in a defect in DNA transfer, but the sequencing analysis was insufficient to identify off target mutations.

### MMJJ_13020 and MMJJ_13030 are essential for transformation and conserved in naturally competent archaea

Three transposon insertions identified in both screens were in MMJJ_13020 and MMJJ_13030, which are predicted to encode an oligonucleotide/oligosaccharide binding (OB-) fold domain protein with a single transmembrane helix and a TctA-like protein with 11 transmembrane helices, respectively. TctA-like proteins are part of the tripartite tricarboxylate transporter (TTT) protein family (32). Because archaea lack identifiable homologs of the bacterial ComEC family of DNA transporters, we hypothesize that another transporter, possibly a TctA-like protein such as MMJJ_13030, is needed for transformation. *M. maripaludis* strains with in-frame deletions of either MMJJ_13020 (Figure 4) or MMJJ_13030 (Figure 5A) were generated and tested for transformation. Both mutant strains were completely defective for transformation, and complementation of the mutation *in trans* rescued this defect.

**Figure 4.**
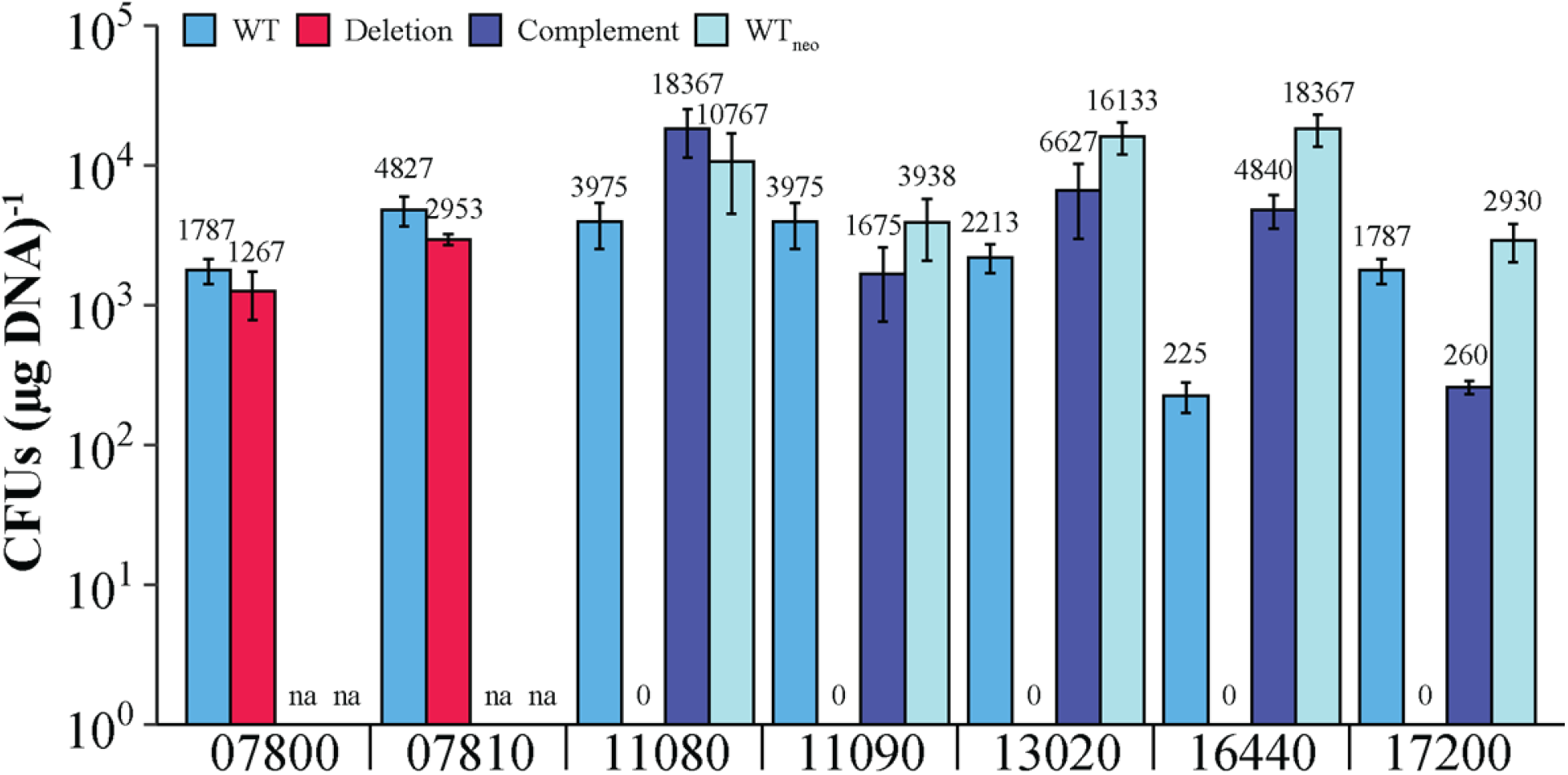
Deletion and complement mutants confirm the role of several candidate genes for transformation: Transformation efficiencies calculated with respect to each of the following gene loci (Genes of interest are referenced using the locus tag format of Poehlein *et. al* (MMJJ_XXXXX)(21)). In each, WT and deletion were performed using pLW40neo as substrate for transformation, while complement and WT_neo_ (WT + pLW40neo – empty vector) were transformed using pLW40. WT and WT_neo_ paired with respective mutants were transformed on the same day with the same transforming DNA. Data are averages from three independent experiments, and error bars represent one standard deviation around the mean.

**Figure 5.**
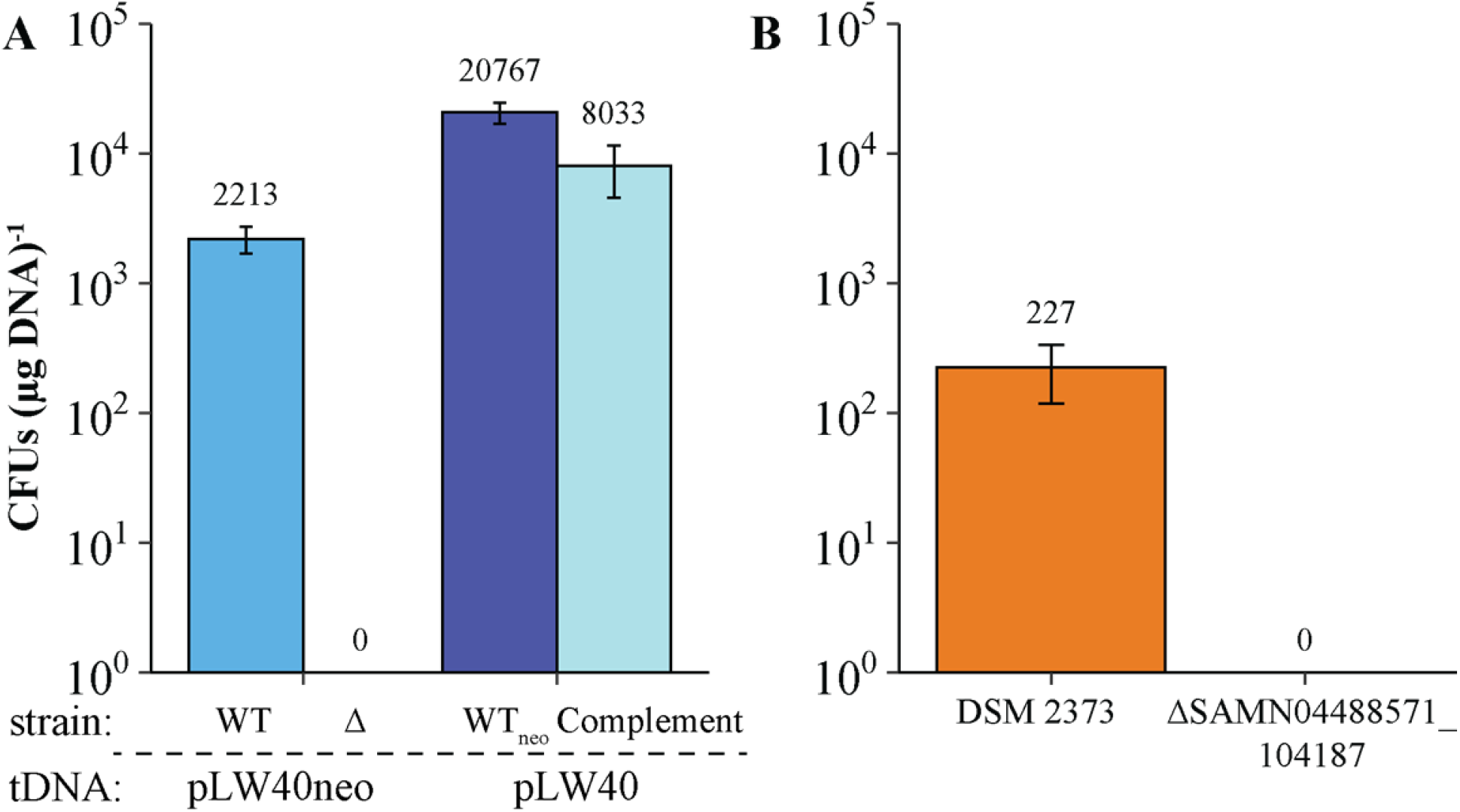
MMJJ_13030 is essential to transformation in *M. maripaludis* and its homolog is essential to transformation in *M. thermophilus* DSM 2373. (A) Transformation efficiencies with and without MMJJ_13030 of *M. maripaludis* JJ. WT_neo_ denotes WT+pLW40neo (empty vector), and complement refers to the Δ*MMJJ_13030*+ pLW40neo-*MMJJ_13030*. WT and WT_neo_ paired with respective mutants were transformed on the same day with the same transforming DNA. Data are averages from three independent experiments, and error bars represent one standard deviation around the mean. (B) Transformation efficiencies of *M. thermophilus* WT (DSM 2373) and a Δ*SAMN04488571_104187* strain transformed on same day with same transforming DNA. Data are averages from three independent experiments, and error bars represent one standard deviation around the mean.

Several components of the natural transformation machinery are likely conserved in naturally competent archaea. For example, the type IV-like pilus is essential for transformation in both *M. maripaludis and M. thermophilus* (14). Because TctA-like proteins are found in several members of the Euryarchaeota, and their function in archaea is unknown, we sought to determine if their role in natural transformation is conserved. From a BLASTp analysis using the standard search criterion/databases, we identified a homolog of MMJJ_13030 in *M. thermophilus* (SAMN04488571_104187); a similar BLASTp search did not retrieve a single clear homolog of MMJJ_13020 in this organism.

We generated an *M. thermophilus* mutant with an in-frame deletion of SAMN04488571_104187 and tested the strain for a transformation defect. Because replicating vectors are not available for *M. thermophilus*, we used the integrating plasmid, pJAL1inter, to test for a transformation defect (14). After incubation with plasmid DNA, we recovered 227 CFUs (µg DNA)^-1^ from wild type *M. thermophilus* on selective media and no transformants from the Δ*SAMN04488571_104187* mutant strain (Figure 5B). Because *M. thermophilus* transformation relies on DNA transfer via natural transformation, and the Δ*SAMN04488571_104187* mutant was completely defective for transformation, we were unable to attempt complementation of this mutation. However, these data suggests that the role of the TctA-like proteins may be functionally conserved in naturally competent archaea.

## Discussion

The ability to acquire DNA by transformation is a major driver of evolution in microbial populations; however, this process is not well understood in archaea. While type IV-like pili are essential to transformation in both bacteria and archaea (14), the remaining proteins required for bacterial transformation are not present in archaea. Here, we employed random mutagenesis to identify genes essential to transformation in *M. maripaludis*. While further work is necessary for validating the roles of these genes, we hypothesize that several play a role in transfer of DNA across the cell membrane. For example, it is unclear whether the archaeal DNA transporter requires ssDNA or dsDNA as a substrate; if ssDNA is required, a secreted thermonuclease like MMJJ_16440 may be important for DNA processing.

MMJJ_13030 is annotated as a TctA-like protein, which is a member of the TTT family of organic acid transporters. This family of proteins is widespread across the Euryarchaeota (32), including other competent archaea such as *Thermococcus kodakarensis, Methanothermobacter marburgensis, Pyrococcus furiosus*, and *Methanococcus voltae* (33–36). In the bacterium *Salmonella typhimurium*, TctABC functions as a citrate transporter (37–41); however, archaea lack clear homologs of TctB and TctC (32). Furthermore, archaeal TctA-like proteins typically encode up to 11 transmembrane helices, in contrast to bacterial proteins that encode up to 12 transmembrane proteins. For these reasons, it is likely that archaeal TctA-like proteins carry out a different function from bacterial TctA. In our screen, MMJJ_13030 was the only membrane transporter identified as important for natural transformation and, along with MMJJ_13020, is predicted to bind DNA as determined by analysis using the DRNApred server (42). A homolog of MMJJ_13030 in *M. thermophilus*, SAMN04488571_104187, is also essential for transformation, suggesting a conserved role for this protein in transformation across multiple naturally competent archaea. While more work needs to be done to determine the exact role in transformation, these features suggest that MMJJ_13030 may encode the archaeal DNA transporter.

We previously showed that type IV-like pili are essential for transformation in both *M. maripaludis* and *M. thermophilus*. Predictably, multiple genes important for expression, processing, and assembly of the type IV-like filament were identified as essential for transformation. Other than pili, histone proteins are essential for transformation *in Thermococcus kodakarensis* (43); however, we did not identify genes encoding histones in our screens so they were not considered further.

Screens for both replicating and integrating vectors identified similar genes important for transformation. pLW40neo is a derivative of pURB500 (27, 44), which is thought to encode the necessary machinery for DNA partitioning. With integrative plasmids like pCRUptNeo recombination into the host chromosome is essential for the retention of selective markers; therefore, we initially hypothesized that additional genes would be essential for this step. In bacterial transformation, recombination can be catalyzed by proteins such as RecA (19, 20). In *M. maripaludis*, there are multiple predicted recombination machineries present, namely RadA (archaeal homolog of RecA/Rad51), and the Mre11-Rad50 double strand break repair system (45). We did not identify either of these systems in our screen, possibly due to overlapping functions of recombination systems present in *M. maripaludis*.

While mapping transposon insertions by sequencing, we found that the *M. maripaludis* strain JJ used in this study lacked ∼80 kb near the gene encoding tRNA^Ser^ (between nucleotides 453,055 and 535,218) that was identified in the reference JJ genome (CP026606 (21)). In the reference genome, this region is flanked by 56 bp of DNA with 100% sequence identity; therefore, it is likely that homologous recombination between these regions resulted in the loss of DNA. Within this 80 kB deletion, 73% of predicted genes (66/89) are annotated as encoding hypothetical proteins or proteins with domains of unknown function. Of the remaining genes, 30% (7/23) appear to be related to transcriptional regulation, suggesting that there may be transcriptional differences between the reference JJ genome and our laboratory strain. Interestingly, this region also contains predicted integrases, a putative restriction endonuclease and associated DNA methyltransferase, and a type II toxin/anti-toxin system. We hypothesize that this region may encode a virus or other mobile genetic element. In any case, the strain of *M. maripaludis* JJ used in this study is naturally competent, so a role for this genomic region was not investigated further.

Despite the fact that several naturally competent archaea have been characterized (14, 46–50), the genes and proteins essential for transformation have not been characterized. Leveraging transposon mutagenesis and protocols for chemical transformation, we have identified several candidate components of the archaeal natural transformation machinery, including pili, DNA binding proteins, nucleases, and predicted transporters. Identification of the genes important for transformation informs efforts to identify potentially competent organisms based on gene content and will broaden understanding of horizontal gene transfer in natural populations.

## Materials and Methods

### Strains, medium, and growth conditions

Strains used in this study are listed in Table 2. *M. maripaludis* strain JJ was acquired from William Whitman and *M. thermophilus* DSM 2373 was purchased from DSMZ (Leibniz Institut, Deutsche Sammlung von Mikroorganismen und Zellkulturen, Braunschweig, Germany).

**Table 2.** Archaeal strains used in this study:

| Strain Name | Genotype | Reference |
| --- | --- | --- |
| KC13 | *M. maripaludis* strain JJΔupt (MMJJ_05990) | (14) |
| KC78 | KC13ΔpstI (MMJJ_06980) | This Study |
| KC120 | KC13ΔMMJJ_07800 | This Study |
| KC87 | KC13ΔMMJJ_07810 | This Study |
| KC103 | KC13ΔMMJJ_11080 | This Study |
| KC104 | KC13ΔMMJJ_11090 | This Study |
| KC123 | KC13ΔMMJJ_13020 | This Study |
| KC122 | KC13ΔMMJJ_13030 | This Study |
| KC89 | KC13ΔMMJJ_16440 | This Study |
| KC121 | KC13ΔMMJJ_17200 | This Study |
| DSM2373 | *M. thermophilus* DSM 2373 WT | (56) |
| KC138 | DSM2373ΔSAMN04488571_104187 | This Study |
| KC19 | KC13+pLW40neo | This Study |
| KC105 | KC103+pLW40neo-MMJJ_11080 | This Study |
| KC106 | KC104+pLW40neo-MMJJ_11090 | This Study |
| KC74 | KC123+pLW40neo-MMJJ_13020 | This Study |
| KC73 | KC122+pLW40neo-MMJJ_13030 | This Study |
| KC107 | KC89+pLW40neo-MMJJ_16440 | This Study |
| KC130 | KC121+pLW40neo-MMJJ_17200 | This Study |

*M. maripaludis* strain JJ and its mutants were grown on McCas medium at 37°C with agitation while *M. thermophilus* DSM 2373 and its mutants were grown in McTry at 55°C as previously described (14). When necessary, the following antibiotics were added to medium at the noted concentrations: neomycin (1 mg mL^−1^ for liquid medium and 0.5 mg mL^−1^ for plates), puromycin (2.5 μg mL^−1^), or 6-azauracil (0.2 mg mL^−1^). *Escherichia coli* DH5α and Rosetta (pLysS) were grown in lysogeny broth or on lysogeny broth 1.5% agar plates supplemented with ampicillin (50 µg mL^-1^) and incubated at 37°C.

### Plasmid construction

Primers are listed in Table 3. All mutants/plasmids were made using the methods described in (14). Briefly, for in-frame genomic deletion mutants in both *M. maripaludis* and *M. thermophilus* 500 bp PCR products of the genomic regions flanking the gene of interest were amplified using primers that encoded 20 bp on either end homologous to pCRUptNeo (51) around the XbaI and NotI restriction sites. Additionally, fragments were constructed to retain the first nine and last twelve nucleotides of the open reading frame. Products were assembled with XbaI/NotI digested pCRUptNeo using NEB builder (# E2621) for Gibson assembly (52). pCRUptNeo has features for propagation in *E. coli* (origin of replication and ampicillin resistance gene) and for selection (neomycin selection) and counterselection (uracil phosphoribosyltransferase) in methanogens. For expression on pLW40neo, genes were PCR amplified with primers that add 20 bp homologous to the regions surrounding AscI and NsiI sites of pLW40neo. These sites place the gene of interest under the control of the *Methanococcus voltae* histone promoter. Assembled constructs were electroporated into *E. coli* DH5α. Transformants were selected on lysogeny broth agar medium containing ampicillin before plasmids were extracted and transferred to methanogens. All constructs were sequence verified by Sanger sequencing at the University of Minnesota Genomics Center or by long Oxford Nanopore through Plasmidsaurus sequencing service (www.plasmidsaurus.com).

**Table 3.**
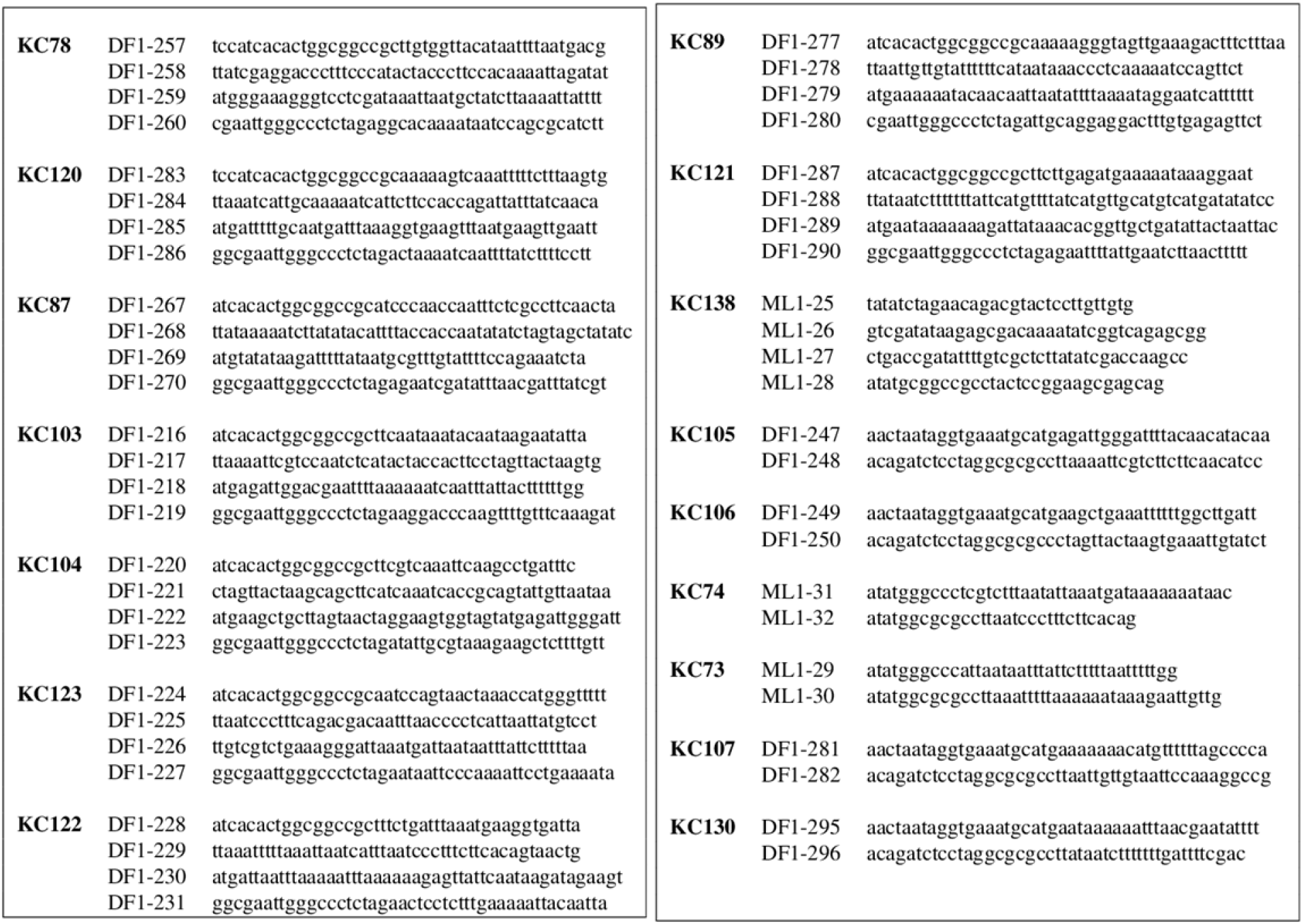
Primers used in this study.

**Table 4.** Plasmid backbones used in this study.

| Name | Description | Reference |
| --- | --- | --- |
| pLW40 | Replicative plasmid in *M. maripaludis*. Contains puromycin resistance cassette. Used as DNA substrate for transformation of complement strains. | (27) |
| pLW40neo | Replicative plasmid in *M. maripaludis*. Contains neomycin resistance cassette. Used as DNA substrate for transformation of deletion mutants/transposon mutants. Also used as a backbone for complementation constructs (under control of *phmv* promoter). | (27) |
| pCRUptNeo | Vector backbone for generating deletion mutants. | (51) |
| pCRUptNeoΔ*flaK* | DNA substrate for testing transformation of integrative vectors. Plasmid integrates either upstream or downstream of the flagella peptidase, *flaK*. | (14) |
| pJJ605 | Non-replicating plasmid (in *M. maripaludis*) carrying mini-mariner transposon. | (22) |
| pT7Tnp | Plasmid carrying HimarI hyperactive transposase expressed under the inducible $\lambda P_L$ promoter | (22) |

### Purification of HimarI

A modified protocol for purification of HimarI from inclusion bodies (22, 53) was performed. Plasmid pT7tnp (22) was transferred into *E. coli* Rosetta (pLysS) via electroporation. Cells were allowed to outgrow for ∼ 30 minutes, then 10 µL was inoculated in 5 mL LB+amp and grown overnight. Five µL was subinoculated into four 250 mL LB with ampicillin added flasks and grown with agitation at 37°C until OD_600_ (Thermo Scientific Genesys 30 spec #840-277000) was between 0.9-1 before expression was induced with 100 μL of 1.25 M IPTG (0.5 mM final concentration) for 3 hours. Cultures were pooled and pelleted by centrifugation at 4000 x g for 10 minutes. Pellets were suspended in 10 mL of resuspension buffer (20mM Tris-HCl (pH 7.6), 25% sucrose, 2mM MgCl_2_, 1mM dithiothreitol (DTT)), distributed into 1 mL aliquots, and flash frozen in liquid nitrogen for long-term storage.

For purification, 2 mL of cell material was thawed at room temperature (RT) then supplemented with 50 μL of 5 mg mL^−1^ lysozyme and incubated at RT for 5 minutes. An equal volume of detergent buffer (2mM Tris-HCl (pH 7.6), 4mM EDTA, 200mM NaCl, 1% Triton X-100, 1% deoxycholate) was added and incubated for 20 minutes at RT. DNase I (75 units) and DNase buffer (final 1x) were added and mixed until the solution was no longer viscous, then incubated for an additional 20 minutes at RT. Pellets containing inclusion bodies were obtained via centrifugation at 16,000 x g for 2 minutes at RT and all remaining steps were performed at 4°C.

Pellets were resuspended with 1 mL wash buffer (0.5% Triton X-100, 1mM EDTA), and pelleted at 16,000 x g for 2 minutes. Pellets were washed an additional 2 times in 1 mL wash buffer, then 2 more times with 1 mL 6M Urea. Pellets were resuspended in 500 μl column buffer (4mM Guanidine HCl, 20mM Tris HCl (pH 7.6), 50mM NaCl, 5mM DTT) and incubated while DEAE-Sepharose Fast Flow (17-0709-10) columns were prepared in column buffer (following manufacturer’s protocol) in disposable columns (Fisher 29924) with a 2-3 mL bed volume. Pellet resuspensions were briefly centrifuged to remove precipitate, then loaded into equilibrated columns. Fifteen 500 µl fractions were collected and visualized via Coomassie staining of an SDS-PAGE gel. Fractions containing transposase were pooled.

Pooled fractions were dialyzed in 10,000 MWCO slide-a-lyzer dialysis cassettes (Thermo scientific #66380) in 500 mL dialysis buffer I (10% glycerol, 25mM Tris HCl (pH 7.6), 50mM NaCl, 2mM DTT, 5mM MgCl_2_) for 5 hours with gentle agitation on an orbital rotating shaker. The cassette was then transferred to 500 mL of dialysis buffer II (10% glycerol, 25mM Tris HCl (pH 7.6), 50mM NaCl, 0.5mM DTT, 5mM MgCl_2_) and gently agitated for an additional 12 hours. Sample was removed then centrifuged at 16,000 x g for 2 minutes to remove precipitate. Supernatant was collected, visualized on an SDS-PAGE gel by Coomassie stain, then the concentration (∼160 µg mL^−1^) was determined via Bradford assay (54) using Coomassie blue reagent (Thermo Scientific #23200). Aliquots (25 µl) were flash frozen in liquid nitrogen and stored at −80°C.

### Optimized transposon mutagenesis

Transposon mutagenesis optimization was based on the methods of Sattler *et al*. (22) and Quitzke *et al*. (23). The following steps were optimized from the original methods: increased concentration of HimarI transposase, elimination of the heat inactivation and ethanol precipitation steps, and a shortened length of incubation. The details of the method are as follows:

On the day of transposon mutagenesis, *M. maripaludis* were subinoculated and grown until an OD_600_ ∼0.7-0.9. While cultures were growing, the transposon reaction was set up. In a total volume of 50 µl, 2x transposon reaction buffer (final concentration of reaction: 25 mM HEPES, 100 mM NaCl, 10 mM MgCl_2_, 2 mM dithiothreitol, pH 7.9), 12.5 µg pJJ605, and 10 µg acylated bovine serum albumin (Promega #R396A) were mixed. Reaction was then initiated with 40-80 µmol of HimarI and incubated at 28°C for 2 hours. Transposon reactions were transferred into *M. maripaludis* using the PEG transformation method (26) with a 4 hour outgrowth. Transformants were selected on McCas + puromycin agar medium and allowed to grow at 37°C for 4 days in anaerobic incubation vessels.

### 96-well natural transformation screen

Following transposon mutagenesis, individual colonies were picked into single wells of 96 well plates (Fisher #12565501) containing 250 µl of non-selective McCas medium. Plates were allowed to incubate at 37°C in anaerobic incubation vessels with 20 mL of 25% Na_2_S for 2 days to ensure wells were grown to stationary phase. Each well was supplemented with 50 µL of McCas mixed with pLW40neo (final concentration 1 µg mL^-1^) and the plates were put back in the anaerobic incubation vessels with 15 mL of fresh 25% Na_2_S and allowed to incubate at 37°C overnight. These plates are referred to as the master plates. 12.5 µl was transferred to one of two replica 96 well plates containing 250 µl McCas + neomycin. Replica plates were performed to remove any false positives that may have arisen from the transfer process. Replica neomycin plates were packaged in anaerobic incubation vessels supplemented with 20 mLs of 25% Na_2_S and allowed to grow at 37°C for 2 days for pLW40neo or 3 days with pCRUptNeoΔ*flaK* while master plates were packaged into anaerobic incubation vessels without any Na_2_S and pressurized to 140 kpa with N_2_:CO_2_ (80:20) at room temperature. Anaerobic incubation vessels were opened and screened for presence or absence of turbidity by eye. Wells where both plates had either no, or unclear growth were marked as candidate competence defective mutants.

Cultures from the master plate that corresponded to the candidate mutants were transferred to 5 mL McCas medium and grown for 24 hours. Any tubes that were not fully grown in 24 hours were marked as slow growth mutants and were excluded from further characterization. Remaining candidates were rescreened for a natural transformation defect (14) with 4 hours of outgrowth. 0.2 mL of outgrowth were inoculated into 5 mL McCas+neomycin and allowed to grow shaking at 37°C. Optical density at 600 nm (OD_600_) was measured at 48 hours. A wild-type culture of *M. maripaludis* strain JJ transformed with 5 µg of pLW40neo typically grows to stationary (OD_600_ ∼1.0) in ∼40 hours with a transformation efficiency average of ∼2000 CFUs (µg DNA)^-1^, so 48 hours was used to try and narrow candidates to only include those essential to transformation. Candidate mutants with an OD_600_>0.2 (limit of detection for growth) at 48 hours were considered false positives, while the rest were prepared for sequencing described below.

### Genomic DNA extractions, sequencing, and mapping

One mL of culture was collected from each mutant and genomic DNA (gDNA) was extracted using Qiagen Blood and Tissue DNeasy kit (#69504). Up to 6 cultures were pooled during the lysis step. Purified gDNA pools were submitted to the Microbial Genome Sequencing center (MiGS) for 2×151bp sequencing using the NextSeq 2000 platform.

For all sequencing alignments, forward sequencing reads were analyzed by local alignment using Bowtie2 (v.2.3.4.1) with default parameters (55). The resulting alignments were visualized using Geneious prime^®^(v.2021.0.3). For identification of transposon insertion loci, reads were first mapped to the plasmid pJJ605 sequence, which contains the transposon. Reads that aligned to pJJ605 were then mapped to the *M. maripaludis* strain JJ genome (Accession CP026606). With this approach, non-transposon reads that corresponded to the *pmcrB* and in the case of integrative plasmids, reads that are shared between pCRUptNeoΔ*flaK*, and pJJ605 were observed. These reads did not share the same insertion pattern as transposon reads, therefore were excluded from analysis. In all samples analyzed, the number of mutants that were pooled equaled the number insertions mapped, validating the sequencing approach.

### Transformation of *M. maripaludis* and *M. thermophilus*

Chemical transformations of *M. maripaludis* were performed as previously described in (26) with the variations described in (14). All natural transformations for both *M. maripaludis* and *M. thermophilus* were performed as described (14).

## Acknowledgements

We thank Michael Rother for providing strains and constructs for the purification of transposases and generation of transposon mutants. This study was supported by the National Science Foundation under grant number MCB-2148165.

## References

1. Wolferen M van, Wagner A, Does C van der, Albers S-V. 2016. The archaeal Ced system imports DNA. Proc Natl Acad Sci USA 113:2496.

2. Greve B, Jensen S, Brügger K, Zillig W, Garrett RA. 2004. Genomic comparison of archaeal conjugative plasmids from Sulfolobus. Archaea 1:231–239.

3. Shalev Y, Turgeman-Grott I, Tamir A, Eichler J, Gophna U. 2017. Cell surface glycosylation is required for efficient mating of Haloferax volcanii. Front Microbiol 8:1253.

4. Prangishvili D. 2013. The wonderful world of archaeal viruses. Ann Rev Microbiol 67:565–585.

5. Villa TG, Feijoo-Siota L, Sánchez-Pérez A, Rama JLR, Sieiro C. 2019. Horizontal gene transfer in bacteria, an overview of the mechanisms involved, p. 3–76. In Villa, TG. Viñas, M (eds.), Horizontal Gene Transfer: Breaking Borders Between Living Kingdoms. Springer International Publishing, Cham.

6. Johnston C, Martin B, Fichant G, Polard P, Claverys J-P. 2014. Bacterial transformation: distribution, shared mechanisms and divergent control. 3. Nat Rev Microbiol 12:181–196.

7. Blokesch M. 2016. Natural competence for transformation. Curr Biol 26:R1126–R1130.

8. Seitz P, Blokesch M. 2013. DNA-uptake machinery of naturally competent Vibrio cholerae. Proc Natl Acad Sci USA 110:17987–17992.

9. Ellison CK, Dalia TN, Ceballos AV, Wang JC-Y, Biais N, Brun YV, Dalia AB. 2018. Retraction of DNA-bound type IV competence pili initiates DNA uptake during natural transformation in Vibrio cholerae. Nat Microbiol 3:773–780.

10. Cehovin A, Simpson PJ, McDowell MA, Brown DR, Noschese R, Pallett M, Brady J, Baldwin GS, Lea SM, Matthews SJ, Pelicic V. 2013. Specific DNA recognition mediated by a type IV pilin. Proc Natl Acad Sci USA 110:3065–3070.

11. Aas FE, Wolfgang M, Frye S, Dunham S, Løvold C, Koomey M. 2002. Competence for natural transformation in Neisseria gonorrhoeae: components of DNA binding and uptake linked to type IV pilus expression. Mol Microbiol 46:749–760.

12. Seitz P, Blokesch M. 2013. Cues and regulatory pathways involved in natural competence and transformation in pathogenic and environmental Gram-negative bacteria. FEMS Microbiol Rev 37:336–363.

13. Corbinais C, Mathieu A, Damke PP, Kortulewski T, Busso D, Prado-Acosta M, Radicella JP, Marsin S. 2017. ComB proteins expression levels determine Helicobacter pylori competence capacity. Sci Rep 7:41495.

14. Fonseca DR, Halim MFA, Holten MP, Costa KC. 2020. Type IV-like pili facilitate transformation in naturally competent archaea. J Bacteriol 202:e00355–20.

15. Piepenbrink KH. 2019. DNA uptake by type IV filaments. Front Mol Biosci 6:1.

16. Angelov A, Bergen P, Nadler F, Hornburg P, Lichev A, Übelacker M, Pachl F, Kuster B, Liebl W. 2015. Novel Flp pilus biogenesis-dependent natural transformation. Front Microbiol 6:84.

17. Pimentel ZT, Zhang Y. 2018. Evolution of the natural transformation protein, ComEC, in bacteria. Front Microbiol 9:2980.

18. Gophna U, Altman-Price N. 2022. Horizontal gene transfer in archaea-from mechanisms to genome evolution. Ann Rev Microbiol 76:481–502.

19. Claverys J-P, Martin B, Polard P. 2009. The genetic transformation machinery: composition, localization, and mechanism. FEMS Microbiol Rev 33:643–656.

20. Kidane D, Ayora S, Sweasy JB, Graumann PL, Alonso JC. 2012. The cell pole: the site of cross talk between the DNA uptake and genetic recombination machinery. Crit Rev Biochem Mol Biol 47:531–555.

21. Poehlein A, Heym D, Quitzke V, Fersch J, Daniel R, Rother M. 2018. Complete genome sequence of the Methanococcus maripaludis Type Strain JJ (DSM 2067), a model for selenoprotein synthesis in archaea. Genome Announcements 6:e00237–18.

22. Sattler C, Wolf S, Fersch J, Goetz S, Rother M. 2013. Random mutagenesis identifies factors involved in formate-dependent growth of the methanogenic archaeon Methanococcus maripaludis. Mol Genet Genomics 288:413–424.

23. Quitzke V, Fersch J, Seyhan D, Rother M. 2018. Selenium-dependent gene expression in Methanococcus maripaludis: Involvement of the transcriptional regulator HrsM. Biochimica et Biophysica Acta (BBA) - General Subjects 1862:2441–2450.

24. Sarmiento F, Mrázek J, Whitman WB. 2013. Genome-scale analysis of gene function in the hydrogenotrophic methanogenic archaeon Methanococcus maripaludis. Proc Natl Acad Sci USA 110:4726–4731.

25. Lampe DJ, Akerley BJ, Rubin EJ, Mekalanos JJ, Robertson HM. 1999. Hyperactive transposase mutants of the Himar1 mariner transposon. Proc Natl Acad Sci USA 96:11428–11433.

26. Tumbula DL, Makula RA, Whitman WB. 1994. Transformation of Methanococcus maripaludis and identification of a Pst I-like restriction system. FEMS Microbiol Lett 121:309–314.

27. Dodsworth JA, Leigh JA. 2006. Regulation of nitrogenase by 2-oxoglutarate-reversible, direct binding of a PII-like nitrogen sensor protein to dinitrogenase. Proc Natl Acad Sci USA 103:9779–9784.

28. Bao J, Mateos E de D, Scheller S. 2022. Efficient CRISPR/Cas12a-based genome editing toolbox for metabolic engineering in Methanococcus maripaludis. bioRxiv https://doi.org/10.1101/2021.12.29.474413.

29. Lutkenhaus J. 2012. The ParA/MinD family puts things in their place. Trends Microbiol 20:411–418.

30. Haft DH, Payne SH, Selengut JD. 2012. Archaeosortases and exosortases are widely distributed systems linking membrane transit with posttranslational modification. J Bacteriol 194:36–48.

31. Abdul Halim MF, Stoltzfus JD, Schulze S, Hippler M, Pohlschroder M. 2017. ArtA-dependent processing of a Tat substrate containing a conserved tripartite structure that is not localized at the C terminus. J Bacteriol 199:e00802–16.

32. Winnen B, Hvorup RN, Saier MH. 2003. The tripartite tricarboxylate transporter (TTT) family. Res Microbiol 154:457–465.

33. Fukui T, Atomi H, Kanai T, Matsumi R, Fujiwara S, Imanaka T. 2005. Complete genome sequence of the hyperthermophilic archaeon Thermococcus kodakaraensis KOD1 and comparison with Pyrococcus genomes. Genome Research 15:352–363.

34. Liesegang H, Kaster A-K, Wiezer A, Goenrich M, Wollherr A, Seedorf H, Gottschalk G, Thauer RK. 2010. Complete genome sequence of Methanothermobacter marburgensis, a methanoarchaeon model organism. J Bacteriol 192:5850–5851.

35. Bridger SL, Lancaster WA, Poole FL, Schut GJ, Adams MWW. 2012. Genome sequencing of a genetically tractable Pyrococcus furiosus strain reveals a highly dynamic genome. J Bacteriol 194:4097–4106.

36. Whitman WB, Woyke T, Klenk H-P, Zhou Y, Lilburn TG, Beck BJ, De Vos P, Vandamme P, Eisen JA, Garrity G, Hugenholtz P, Kyrpides NC. 2015. Genomic encyclopedia of bacterial and archaeal type strains, phase III: the genomes of soil and plant-associated and newly described type strains. Standards in Genomic Sciences 10:26.

37. Ashton DM, Sweet GD, Somers JM, Kay WW. 1980. Citrate transport in Salmonella typhimurium: studies with 2-fluoro-L-erythro-citrate as a substrate. Can J Biochem Physiol 58:797–803.

38. Somers JM, Sweet GD, Kay WW. 1981. Flurorcitrate resistant tricarboxylate transport mutants of Salmonella typhimurium. Mol Genet Genomics 181:338–345.

39. Sweet GD, Somers JM, Kay WW. 1979. Purification and properties of a citrate-binding transport component, the C protein of Salmonella typhimurium. Can J Biochem Physiol 57:710–715.

40. Sweet GD, Kay CM, Kay WW. 1984. Tricarboxylate-binding proteins of Salmonella typhimurium. Purification, crystallization, and physical properties. J Biol Chem 259:1586–1592.

41. Widenhorn KA, Somers JM, Kay WW. 1988. Expression of the divergent tricarboxylate transport operon (tctI) of Salmonella typhimurium. J Bacteriol 170:3223–3227.

42. Yan J, Kurgan L. 2017. DRNApred, fast sequence-based method that accurately predicts and discriminates DNA-and RNA-binding residues. Nuc Acids Res 45:e84.

43. Čuboňová L, Katano M, Kanai T, Atomi H, Reeve JN, Santangelo TJ. 2012. An archaeal histone is required for transformation of Thermococcus kodakarensis. J Bacteriol 194:6864–6874.

44. Tumbula DL, Bowen TL, Whitman WB. 1997. Characterization of pURB500 from the archaeon Methanococcus maripaludis and construction of a shuttle vector. J Bacteriol 179:2976–2986.

45. Hendrickson EL, Kaul R, Zhou Y, Bovee D, Chapman P, Chung J, Conway de Macario E, Dodsworth JA, Gillett W, Graham DE, Hackett M, Haydock AK, Kang A, Land ML, Levy R, Lie TJ, Major TA, Moore BC, Porat I, Palmeiri A, Rouse G, Saenphimmachak C, Söll D, Van Dien S, Wang T, Whitman WB, Xia Q, Zhang Y, Larimer FW, Olson MV, Leigh JA. 2004. Complete genome sequence of the genetically tractable hydrogenotrophic methanogen Methanococcus maripaludis. J Bacteriol 186:6956–6969.

46. Sato T, Fukui T, Atomi H, Imanaka T. 2003. Targeted gene disruption by homologous recombination in the hyperthermophilic archaeon Thermococcus kodakaraensis KOD1. J Bacteriol 185:210–220.

47. Worrell VE, Nagle DP, McCarthy D, Eisenbraun A. 1988. Genetic transformation system in the archaebacterium Methanobacterium thermoautotrophicum Marburg. J Bacteriol 170:653–656.

48. Lipscomb GL, Stirrett K, Schut GJ, Yang F, Jenney FE, Scott RA, Adams MWW, Westpheling J. 2011. Natural competence in the hyperthermophilic archaeon Pyrococcus furiosus facilitates genetic manipulation: construction of markerless deletions of genes encoding the two cytoplasmic hydrogenases. App Environ Microbiol 77:2232–2238.

49. Patel GB, Nash JHE, Agnew BJ, Sprott GD. 1994. Natural and electroporation-mediated transformation of Methanococcus voltae protoplasts. App Environ Microbiol 60:903–907.

50. Wagner A, Whitaker RJ, Krause DJ, Heilers J-H, van Wolferen M, van der Does C, Albers S-V. 2017. Mechanisms of gene flow in archaea. Nat Rev Microbiol 15:492–501.

51. Costa KC, Wong PM, Wang T, Lie TJ, Dodsworth JA, Swanson I, Burn JA, Hackett M, Leigh JA. 2010. Protein complexing in a methanogen suggests electron bifurcation and electron delivery from formate to heterodisulfide reductase. Proc Natl Acad Sci USA 107:11050–11055.

52. Gibson DG, Young L, Chuang R-Y, Venter JC, Hutchison CA, Smith HO. 2009. Enzymatic assembly of DNA molecules up to several hundred kilobases. Nat Methods 6:343–345.

53. Lampe DJ, Churchill ME, Robertson HM. 1996. A purified mariner transposase is sufficient to mediate transposition in vitro. EMBO J 15:5470–5479.

54. Bradford MM. 1976. A rapid and sensitive method for the quantitation of microgram quantities of protein utilizing the principle of protein-dye binding. Anal Biochem 72:248–254.

55. Langmead B, Salzberg SL. 2012. Fast gapped-read alignment with Bowtie 2. Nat Meth 9:357–359.

56. Rivard CJ, Smith PH. 1982. Isolation and characterization of a thermophilic marine methanogenic bacterium, Methanogenium thermophilicum sp. nov.†. Int J Syst Evol Microbiol, 32:430–436.

